# DropletFactory CORE – a droplet cytometry and sorting platform for fast and accessible screening in biotechnology

**DOI:** 10.64898/2026.03.11.711014

**Authors:** Robin Veere, Merle N. Zenner, Anum Afroz, Rauno Jõemaa, Triini Olman, Simona Bartkova, Steven A. van der Hoek, Azra Melkic, Alice J.-L. Zheng, András J. Laki, Mária Laki, Tamás Pardy, Ott Scheler

**Affiliations:** Department of Chemistry and Biotechnology, Tallinn University of Technology, Tallinn, Estonia; Thomas Johann Seebeck Department of Electronics, Tallinn University of Technology, Tallinn, Estonia; TFTAK AS, Tallinn, Estonia; Institute of Technology, University of Tartu, Tartu, Estonia; Faculty of IT and Bionics, Pazmany Peter Catholic University, Budapest, Hungary

**Keywords:** Droplet microfluidics, droplet cytometry, droplet sorting, cell screening, biotechnology instrumentation

## Abstract

Droplet sorting technology has the potential to revolutionize the biotechnology sector as it provides massive high-throughput screening capacity, but the technology remains not accessible for a wider audience yet. There is a need for more affordable droplet sorting platforms to design cell factories and screen cell libraries. In here we demonstrate our droplet cytometry/sorter platform for single-cell screening of yeast cells based on their fluorescence signal.

## 1. Introduction

### 1.1. Current droplet sorting technologies

Fluorescence-activated droplet sorting (FADS) represents the current state-of-the-art method for high-throughput selection of picolitre to nanolitre sized droplets, combining optical screening with real-time classification and rapid sorting. In typical systems, droplets are excited by a focused laser beam, and the resulting fluorescence signal is detected by a sensitive photomultiplier tube (PMT). When a fluorescence signal exceeds a predefined threshold, precisely timed electric field pulses redirect selected droplets using dielectrophoretic forces.

Early implementations demonstrated reliable sorting at kilohertz frequencies for enzyme and cell-based screening applications [1], [2]. Following recent advances in microfluidic chip design, timing electronics and real-time signal processing have further increased throughput, stability and sorting accuracy [3]. Modern platforms also enable multiplexed sorting and automated feedback control, improving robustness during long-term screening experiments [4], [5].

Apart from dielectrophoretic approaches, alternative active redirection strategies such as acoustophoretic and magnetophoretic droplet sorting have emerged, offering improved biocompatibility for sensitive biological samples [6], [7]. Despite these technological developments, droplet sorting platforms often remain complex systems that require precise optical alignment, specialized hardware and significant operational expertise [8].

As a result, many implementations remain limited to specialized laboratories, narrowing broader adoption of droplet microfluidics sorting in biotechnology. These limitations motivate the development of simpler and more scalable droplet analysis platforms that maintain high analytical performance while improving accessibility and adoption.

### 1.2. Applications in biotechnology

Droplet sorting technologies have emerged as a cornerstone of modern biotechnology by enabling ultrahigh-throughput, compartmentalized analysis and selection of single cells or microcolonies. These platforms combine droplet generation, incubation, detection and sorting to maintain genotype-phenotype linkage within individual microreactors, enabling selection across millions of independent experiments.

As a result, droplet-based screening has accelerated applications such as directed enzyme evolution, biocatalyst and enzyme discovery, microbial ecology studies, and optimization of engineered cell factories [9], [10], [11], [12], [13].

Foundational work established the engineering principles required for high-speed droplet classification and phenotypic selection across diverse biological system. More recent studies have focused on improving data quality and functional resolution by combining FADS into multi-step workflows that enrich high-quality single cell encapsulations and improve signal-to-noise ratio [14].

In parallel, complementary developments have introduced passive or simplified droplet analysis strategies that reduce instrumentation requirements, highlighting the growing interest in scalable and accessible microfluidic screening technologies [15]. Together, these advances reflect a progression toward modular, automated and information-rich droplet sorting platforms capable of supporting increasingly complex biological screening workflows.

### 1.3. Goal of this work

This work presents an early technical preview of the DropletFactory CORE FADS platform, integrating optical fluorescence detection, real-time signal processing and dielectrophoretic droplet redirection within a modular experimental setup designed for flexible droplet analysis and sorting.

System performance was evaluated by characterizing optical detection sensitivity using fluorescent dye droplets, analysing fluorescence distributions from GFP expressing yeast droplets, and performing proof-of-concept sorting experiments. These studies demonstrate the operational principles and functionality of the platform for fluorescence-based droplet analysis.

This early preview of the DropletFactory CORE system aims to disclose the high-level system design, sensitivity evaluation with dye-filled droplets, as well as an early proof-of-concept evaluation on droplet-assisted yeast sorting.

## 2. Methodology

### 2.1. Sorting system

FADS was implemented through an integrated microfluidic, optical and electronic workflow (Figure 1), in which droplets are detected, classified and selectively redirected using short electric field pulses.

**Figure 1.**
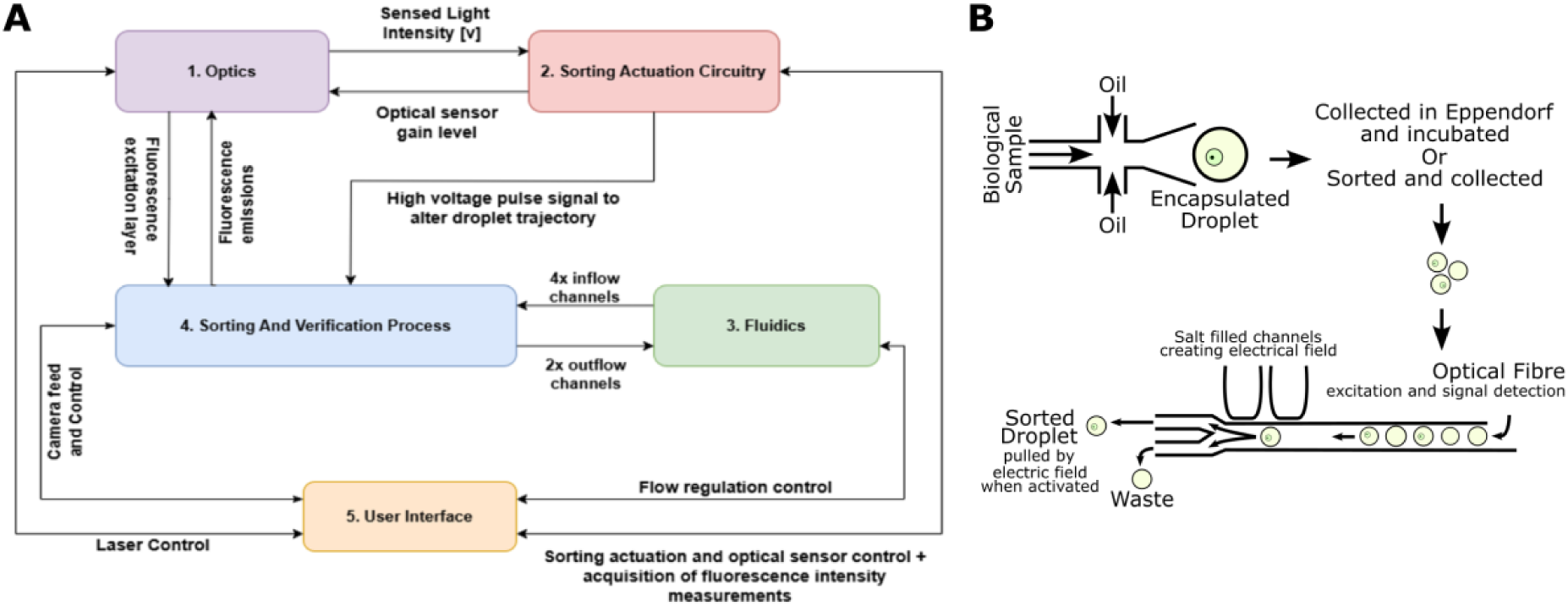
Overview of DropletFactoy CORE system workflow. (A) Block diagram of the sorting system. (B) Schematic of sorting experiment steps and workflow.

The DropletFactory CORE experimental platform is a modification of previously described FADS systems [1], [14] and follows the established principle of optical fluorescence detection coupled with electric field induced dielectrophoretic droplet sorting.

Droplets were generated prior to sorting using a flow-focusing microfluidic chip. The continuous phase consisted of HFE 7500 fluorocarbon oil (3M™ Novec™) supplemented with 2% (w/w) poly(ethylene glycol)-perfluoropolyether (PEG-PFPE) triblock surfactant [16]. The dispersed phase depended on the specific experiment and consisted either of a fluorescent dye-water mixture or of growth medium containing yeast cells. Gastight syringes (Hamilton ®) were operated using NE-501 syringe pumps (New Era Pump Systems, Inc) with custom software (Figure 2).

**Figure 2.**
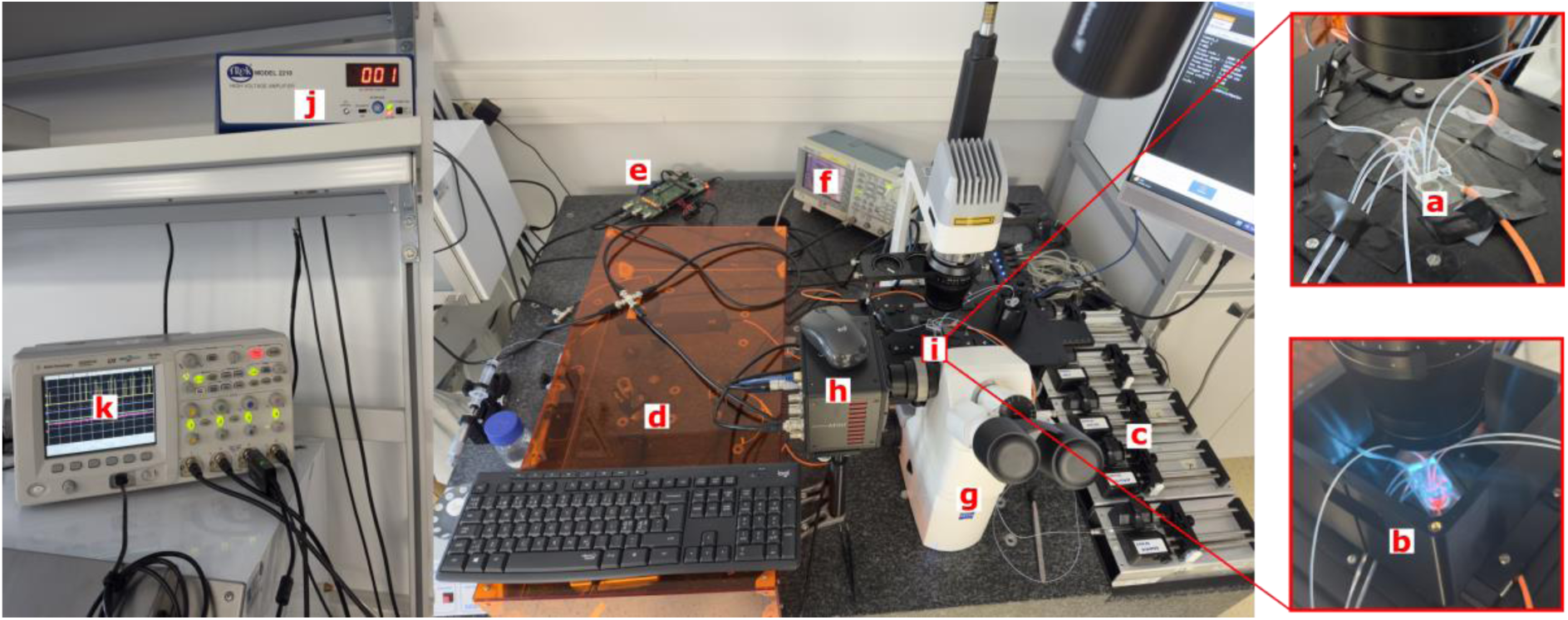
Setup of DropletFacory CORE sorting apparatus. Sorting apparatus consists of microfluidic sorting chip (a), 3D-printed microfluidic chip box (b), syringe pumps (c), optical components (lasers, filters, photomultiplier tube) under protective housing (d), signal acquisition and processing system (e), function generator (f), inverted microscope (g), high-speed camera (h), microfluidic chip holder (i), voltage amplifier (j), and oscilloscope (k). Main system components and manufacturers summarized in Appendix Table 1.

**Table 1.**
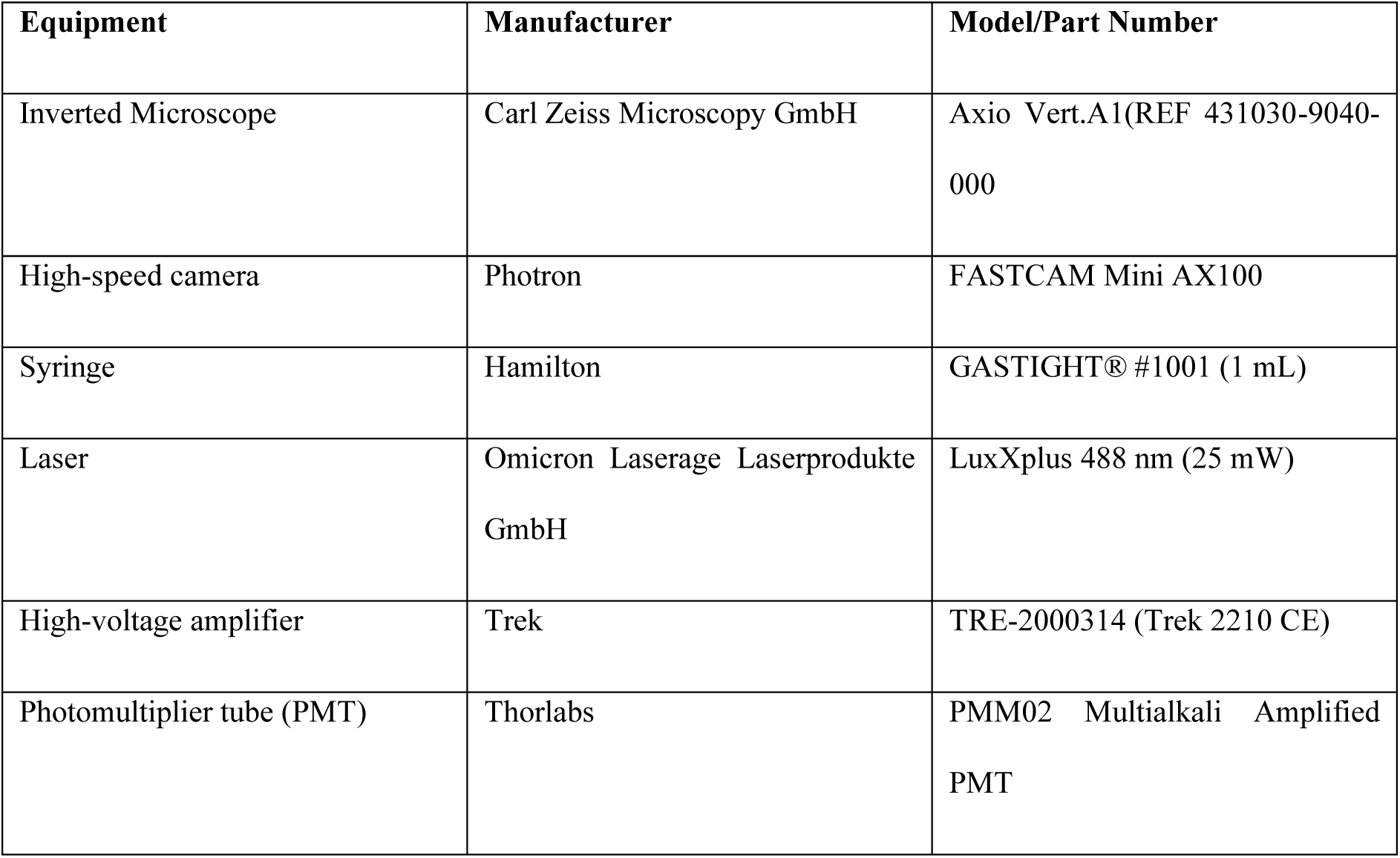
Main system components and manufacturers.

Yeast cells were grown overnight from single colonies and diluted to the desired concentration, resulting in approximately 38% of the generated droplets being empty, around 37% to contain a single cell, and about 25% to contain two or more cells. Droplets were collected in Eppendorf tubes and incubated on the lab bench overnight (approximately 16 h from generation to the start of the sorting experiment)

Droplet-in-oil samples were loaded into tubing connected to a syringe and inserted into the corresponding inlet of the droplet sorting chip (Figure 2). The microfluidic device was placed inside an enclosed chip box (Figure 2) under dark conditions to minimize background light and prevent interference with the optical detection system.

Within the sorting chip, droplets were spaced and passed regularly through the optical detection and sorting region. Flow rates and pressures inside the device were allowed to stabilize until a stable droplet frequency was achieved, after which sorting commenced.

A 488 nm excitation laser was coupled into the detection region via an optical fibre to excite fluorescent cells within the passing droplets. Emitted fluorescence was collected and directed to a PMT (Figure 2), which converted photon flux (light intensity over time) into an electrical signal proportional to fluorescence intensity. Fluorescence signals were recorded at high throughput and processed in real time using a Raspberry Pi-based data acquisition system (Figure 2).

A fluorescence intensity threshold was defined based on the recorded signal distribution. When a droplet generated signal exceeded this threshold, a trigger pulse was transmitted to a function generator and voltage amplifier (Figure 2), producing an alternating current square-wave signal. Milliseconds long voltage pulse was applied to the sorting electrodes, which were filled with 5 M NaCl solution, after a predefined delay corresponding to the droplet travel time between the detection region and the sorting junction

Application of the electric field generated a dielectrophoretic force that redirected droplets containing fluorescent cells into the designated collection outlet. Droplets that did not exceed the threshold remained on the default flow path and were transported to the waste outlet by a continuous bias flow applied through a dedicated bias channel.

The pulse duration, delay and applied voltage were adjustable parameters optimized according to droplet velocity, flow rate and desired sorting throughput. The entire process was visually monitored using a microscope, high-speed camera and an oscilloscope (Figure 2).

### 2.2. GFP expressing yeast

#### 2.2.1. Strains and media composition

*Sc-*SpCas9, a yeast strain expressing SpCas9 derived from *Saccharomyces cerevisiae* CEN.PK113-7D was used as the parental strain for obtaining GFP-expressing strains. *Escherichia coli* DH5ɑ was used for molecular cloning.

Yeast strains were cultured at 30℃ and 250 rpm in either yeast extract peptone dextrose (YPD) or chemically defined medium (CDM). The composition of both cultivation media can be found in Table 2. Pre-cultures were grown overnight (16-18 h) from single colonies. For transformations, pre-cultures were reinoculated in 50 mL YPD to reach OD_600_ ≈ 2 after overnight incubation. For sorter experiments, yeasts were grown overnight in 5 mL CDM and diluted to the desired cell concentration. Agar plates contained 20 g/L agar and were incubated at 30℃ (*S, cerevisiae*) or 37℃ (*E. coli*).

**Table 2.** Medium for yeasts and bacteria.

| Medium | Component | Concentration |
| --- | --- | --- |
| YPD | Yeast extract | 10 g/L |
|  | Peptone | 20 g/L |
|  | Glucose | 20 g/L |
| Chemically defined media (CDM) | Glucose | 20 g/L |
|  | (NH<sub>4</sub>)<sub>2</sub>SO<sub>4</sub> | 5 g/L |
|  | KH<sub>2</sub>PO<sub>4</sub> | 3 g/L |
|  | MgSO<sub>4</sub>·7H<sub>2</sub>O | 500 mg/L |
|  | Trace element solution | 1 mL/L |
|  | Vitamin solution | 0.5 mL/L |
| Trace element solution | FeSO<sub>4</sub>·7H<sub>2</sub>O | 3 mg/L |
|  | EDTA | 15 mg/L |
|  | H<sub>3</sub>BO<sub>3</sub> | 1 mg/L |
|  | Na<sub>2</sub>MoO<sub>4</sub>·4H<sub>2</sub>O | 0.4 mg/L |
|  | KI | 0.1 mg/L |
|  | CaCl<sub>2</sub>·2H<sub>2</sub>O | 4.5 mg/L |
|  | ZnSO<sub>4</sub>·7H<sub>2</sub>O | 4.5 mg/L |
|  | MnCl<sub>2</sub>·4H<sub>2</sub>O | 1 mg/L |
|  | CoCl<sub>2</sub>·6H<sub>2</sub>O | 0.3 mg/L |
|  | CuSO<sub>4</sub>·5H<sub>2</sub>O | 0.3 mg/L |
| Vitamin solution | Biotin | 0.05 mg/L |
|  | 4-Aminobenzoic acid | 0.2 mg/L |
|  | Nicotinic acid | 1 mg/L |
|  | Calcium pantothenate | 1 mg/L |
|  | Pyridoxine hydrochloride | 1 mg/L |
|  | Thiamine hydrochloride | 1 mg/L |
|  | Myo-inositol | 0.025 mg/L |
| LB | Yeast extract | 5 g/L |
|  | Tryptone | 10 g/L |
|  | NaCl | 10 g/L |

#### 2.2.2. Plasmid construction and yeast transformation

Plasmids were constructed according to the Golden Gate assembly protocol of the Yeast Modular Cloning (MoClo) toolkit [17]. Instead of the Yeast MoClo entry vector, an adapted entry vector using kanamycin resistance and BpiI as the Type IIS restriction enzyme was used. Promoter sequences of *S. cerevisiae* were PCR-amplified (Phusion Hot Start II High-Fidelity PCR Master Mix, Thermo Scientific) from *S. cerevisiae* genomic DNA or ordered as synthetic DNA (Twist Bioscience), summarized in Table 5. The oligos used for PCR amplification and colony PCR (cPCR) are described in Table 6. The DNA fragments were cloned into the entry vector using FastDigest BpiI (Thermo Scientific), forming part plasmids. Part plasmids (Table 4) carrying promoters, GFP, and terminators were subsequently assembled into a destination plasmid targeting the X-2 integration site of the EasyClone MarkerFree method [17] using FastDigest Eco31I (Thermo Scientific) to form the final plasmids (Table 4).

**Table 3.**
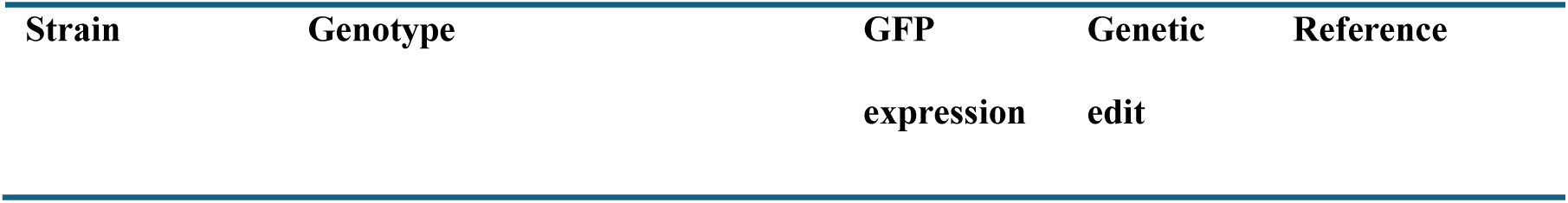
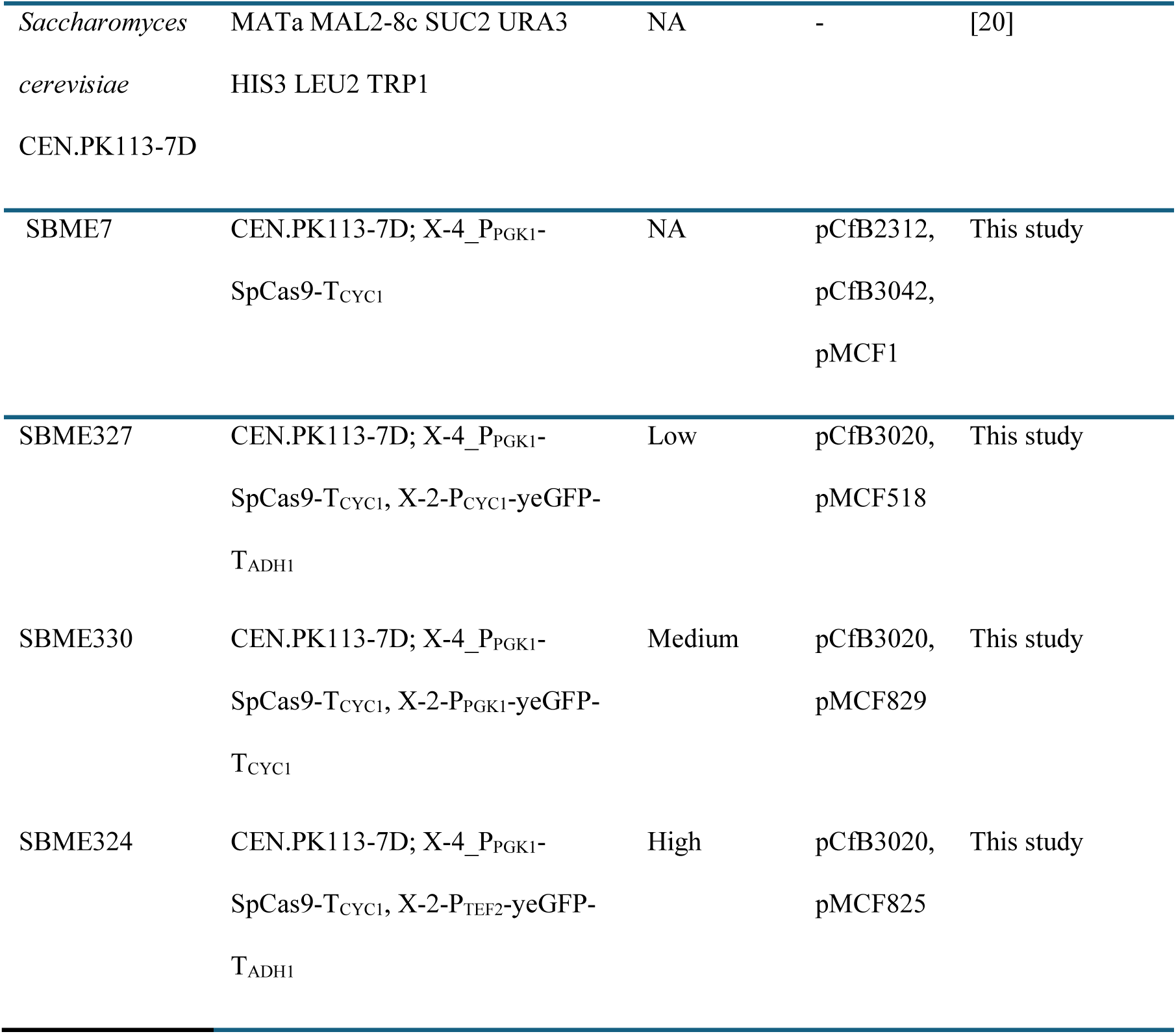
Yeast strains used in this study.

**Table 4.**
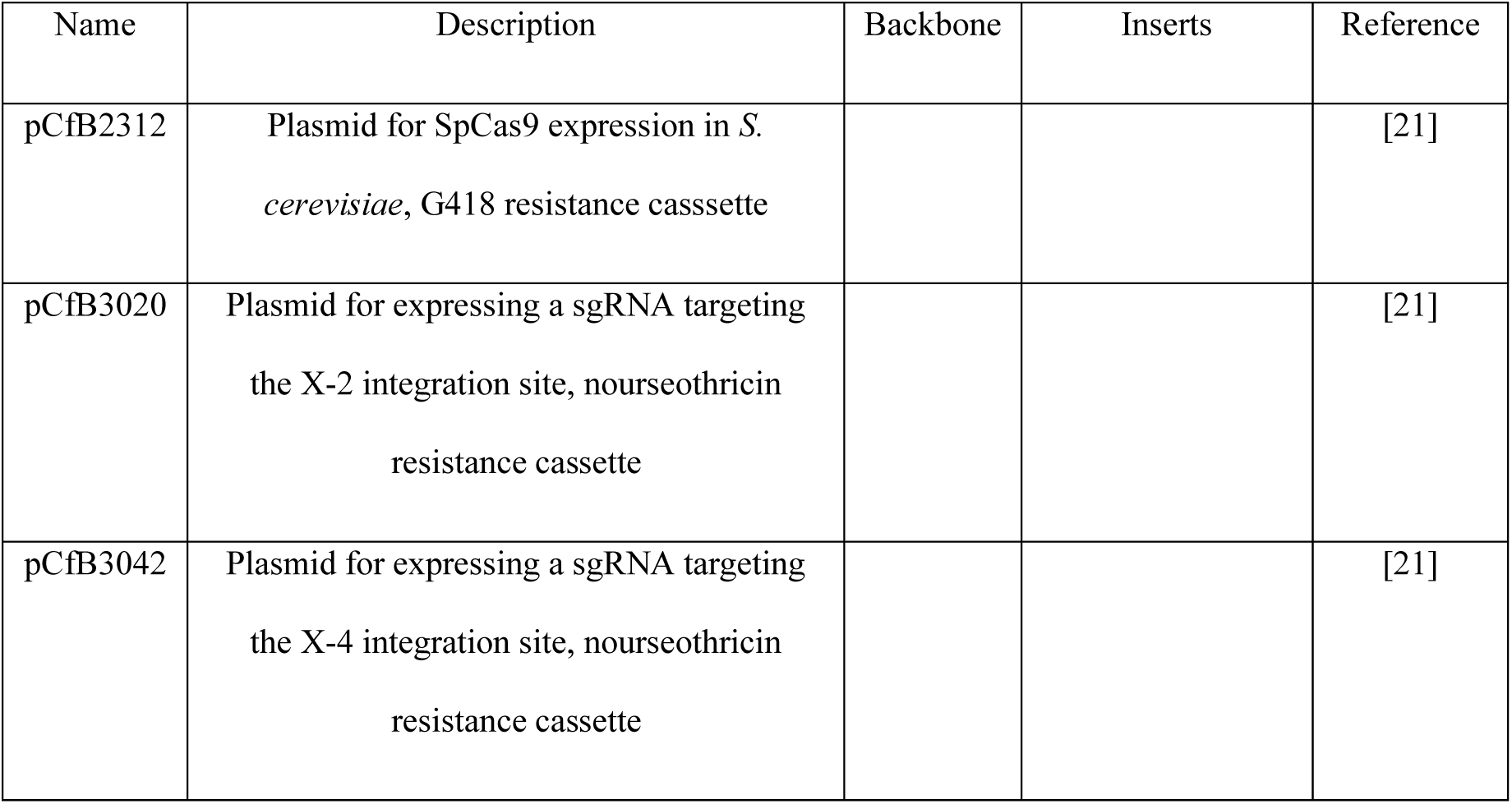
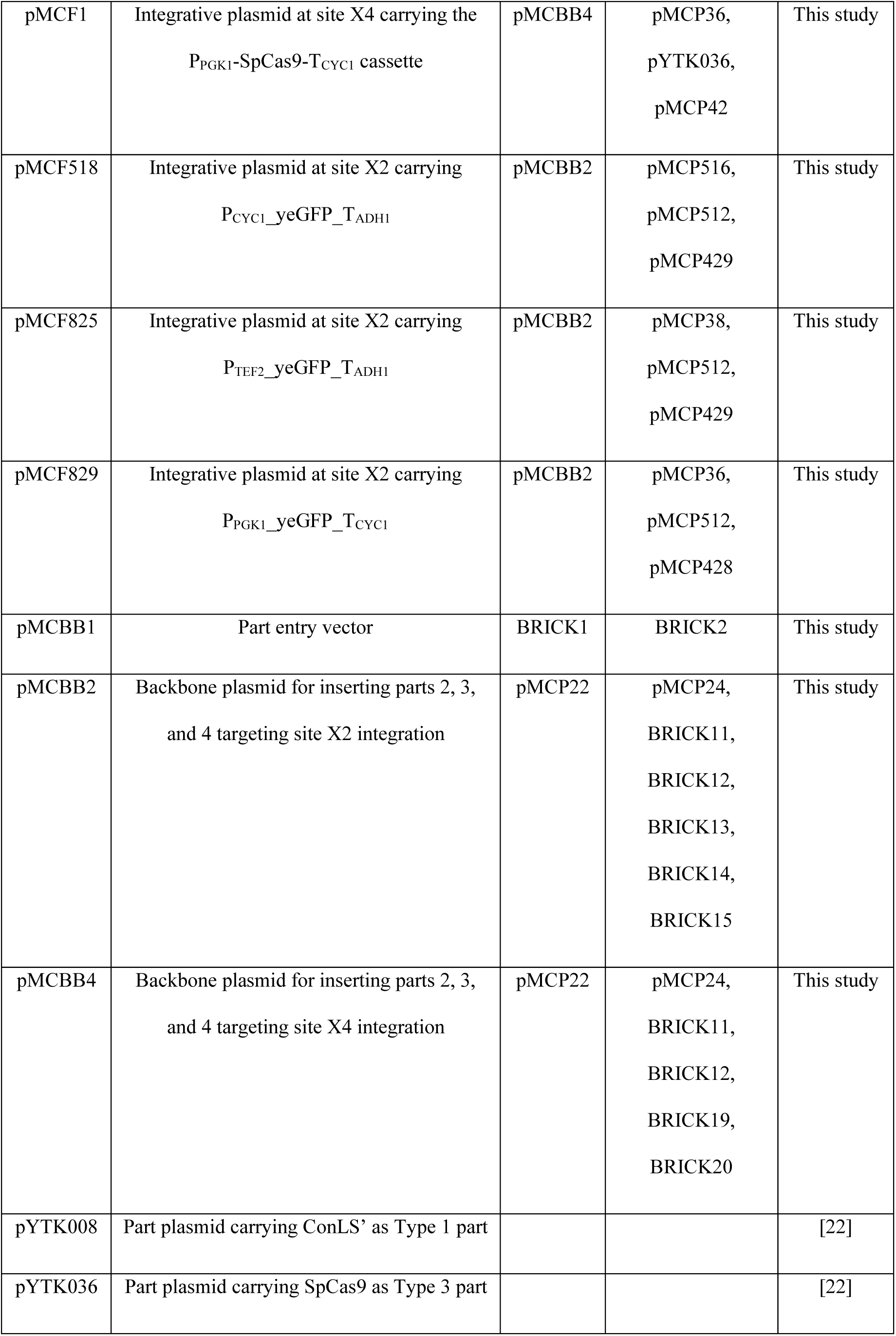
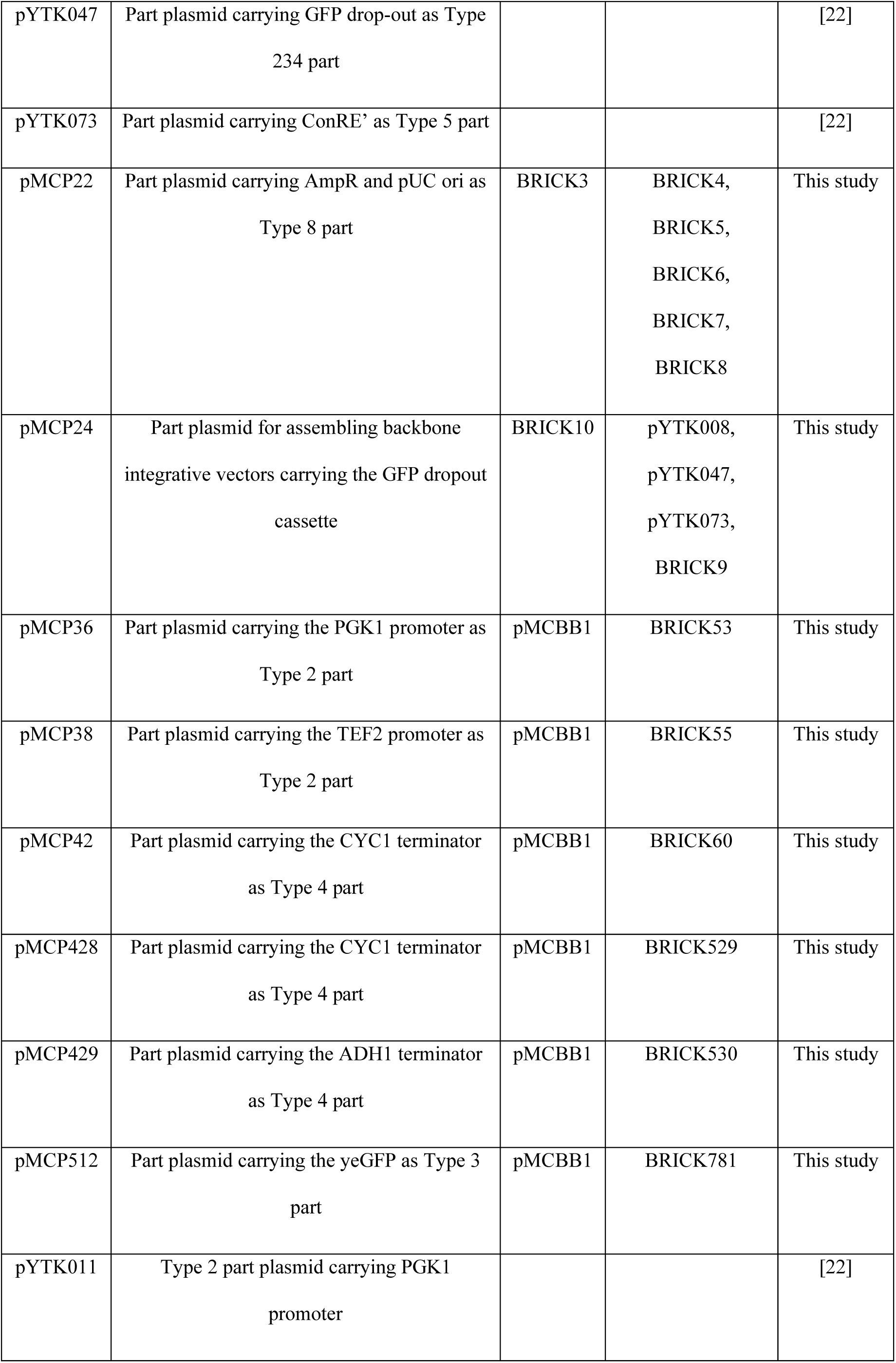
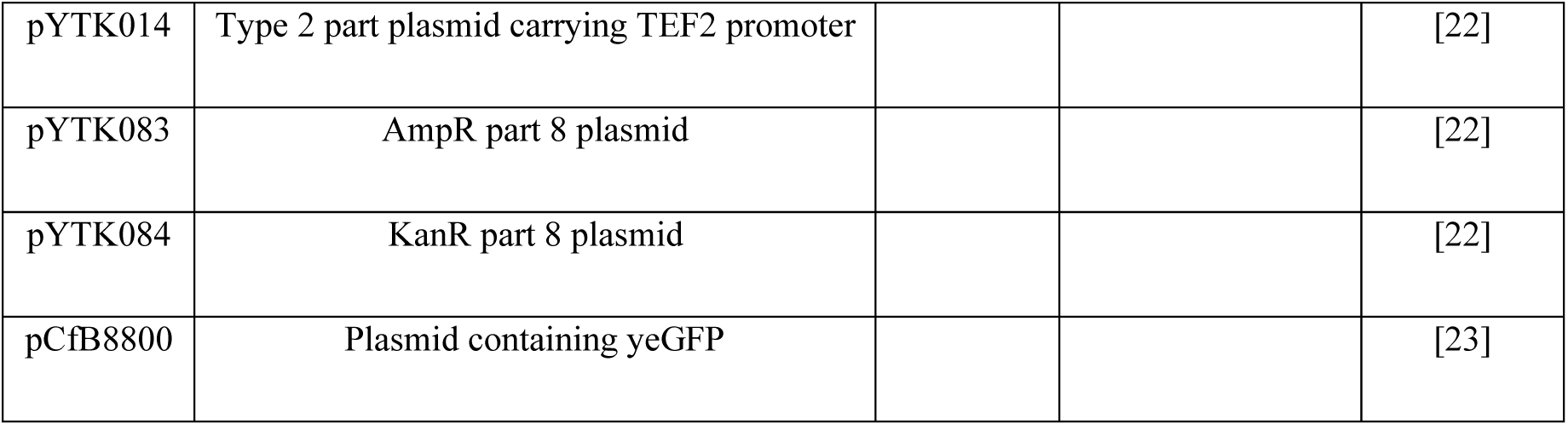
List of plasmids used in this study.

| Name | Description | Backbone | Inserts | Reference |
| --- | --- | --- | --- | --- |
| pCfB2312 | Plasmid for SpCas9 expression in *S. cerevisiae*, G418 resistance cassette |  |  | [21] |
| pCfB3020 | Plasmid for expressing a sgRNA targeting the X-2 integration site, nourseothricin resistance cassette |  |  | [21] |
| pCfB3042 | Plasmid for expressing a sgRNA targeting the X-4 integration site, nourseothricin resistance cassette |  |  | [21] |
| pMCF1 | Integrative plasmid at site X4 carrying the<br>P<sub>PGK1</sub>-SpCas9-T<sub>CYC1</sub> cassette | pMCBB4 | pMCP36,<br>pYTK036,<br>pMCP42 | This study |
| pMCF518 | Integrative plasmid at site X2 carrying<br>P<sub>CYC1</sub>_yeGFP_T<sub>ADH1</sub> | pMCBB2 | pMCP516,<br>pMCP512,<br>pMCP429 | This study |
| pMCF825 | Integrative plasmid at site X2 carrying<br>P<sub>TEF2</sub>_yeGFP_T<sub>ADH1</sub> | pMCBB2 | pMCP38,<br>pMCP512,<br>pMCP429 | This study |
| pMCF829 | Integrative plasmid at site X2 carrying<br>P<sub>PGK1</sub>_yeGFP_T<sub>CYC1</sub> | pMCBB2 | pMCP36,<br>pMCP512,<br>pMCP428 | This study |
| pMCBB1 | Part entry vector | BRICK1 | BRICK2 | This study |
| pMCBB2 | Backbone plasmid for inserting parts 2, 3,<br>and 4 targeting site X2 integration | pMCP22 | pMCP24,<br>BRICK11,<br>BRICK12,<br>BRICK13,<br>BRICK14,<br>BRICK15 | This study |
| pMCBB4 | Backbone plasmid for inserting parts 2, 3,<br>and 4 targeting site X4 integration | pMCP22 | pMCP24,<br>BRICK11,<br>BRICK12,<br>BRICK19,<br>BRICK20 | This study |
| pYTK008 | Part plasmid carrying ConLS' as Type 1 part |  |  | [22] |
| pYTK036 | Part plasmid carrying SpCas9 as Type 3 part |  |  | [22] |
| pYTK047 | Part plasmid carrying GFP drop-out as Type 234 part |  |  | [22] |
| pYTK073 | Part plasmid carrying ConRE' as Type 5 part |  |  | [22] |
| pMCP22 | Part plasmid carrying AmpR and pUC ori as Type 8 part | BRICK3 | BRICK4,<br>BRICK5,<br>BRICK6,<br>BRICK7,<br>BRICK8 | This study |
| pMCP24 | Part plasmid for assembling backbone integrative vectors carrying the GFP dropout cassette | BRICK10 | pYTK008,<br>pYTK047,<br>pYTK073,<br>BRICK9 | This study |
| pMCP36 | Part plasmid carrying the PGK1 promoter as Type 2 part | pMCBB1 | BRICK53 | This study |
| pMCP38 | Part plasmid carrying the TEF2 promoter as Type 2 part | pMCBB1 | BRICK55 | This study |
| pMCP42 | Part plasmid carrying the CYC1 terminator as Type 4 part | pMCBB1 | BRICK60 | This study |
| pMCP428 | Part plasmid carrying the CYC1 terminator as Type 4 part | pMCBB1 | BRICK529 | This study |
| pMCP429 | Part plasmid carrying the ADH1 terminator as Type 4 part | pMCBB1 | BRICK530 | This study |
| pMCP512 | Part plasmid carrying the yeGFP as Type 3 part | pMCBB1 | BRICK781 | This study |
| pYTK011 | Type 2 part plasmid carrying PGK1 promoter |  |  | [22] |
| pYTK014 | Type 2 part plasmid carrying TEF2 promoter |  |  | [22] |
| pYTK083 | AmpR part 8 plasmid |  |  | [22] |
| pYTK084 | KanR part 8 plasmid |  |  | [22] |
| pCfB8800 | Plasmid containing yeGFP |  |  | [23] |

**Table 5.** List of DNA fragments used in this study.

| Name | Template | Oligos | Description |
| --- | --- | --- | --- |
| BRICK1 | pYTK084 | OL1, OL2 | KanR-ColE1 amplification |
| BRICK2 | pYTK047 | OL3, OL4 | GFP drop-out cassette with BpiI sites amplification |
| BRICK3 | pYTK083 | OL5, OL6 | AmpR-ColE1 amplification |
| BRICK4 | pYTK083 | OL7, OL8 | Part 1 of RFP drop-out cassette amplification to remove BpiI sites |
| BRICK5 | pYTK083 | OL9, OL10 | Part 2 of RFP drop-out cassette amplification to remove BpiI sites |
| BRICK6 | pYTK083 | OL11, OL12 | Part 3 of RFP drop-out cassette amplification to remove BpiI sites |
| BRICK7 | pYTK083 | OL13, OL14 | Part 4 of RFP drop-out cassette amplification to remove BpiI sites |
| BRICK8 | pYTK083 | OL15, OL16 | Part 5 of RFP drop-out cassette amplification to remove BpiI sites |
| BRICK9 |  | OL17, OL18 | Spacer amplification |
| BRICK10 | pYTK084 | OL19, OL20 | KanR-ColE1 amplification |
| BRICK11 |  | OL21, OL22 | Annealing DNA for standardized cPCR location on plasmid |
| BRICK12 |  | OL23, OL24 | Annealing DNA for standardized cPCR<br>location on plasmid |
| BRICK13 | *S. cerevisiae* genomic DNA | OL25, OL26 | X-2 UP site amplification |
| BRICK14 | *S. cerevisiae* genomic DNA | OL27, OL28 | Part 1 of X-2 DOWN site amplification |
| BRICK15 | *S. cerevisiae* genomic DNA | OL29, OL30 | Part 2 of X-2 DOWN site amplification |
| BRICK19 | *S. cerevisiae* genomic DNA | OL37, OL38 | X-4 UP site amplification |
| BRICK20 | *S. cerevisiae* genomic DNA | OL39, OL40 | X-4 DOWN site amplification |
| BRICK53 | pYTK011 | OL133, OL134 | PGK1 promoter amplification |
| BRICK55 | pYTK014 | OL137, OL138 | TEF2 promoter amplification |
| BRICK60 | *S. cerevisiae* genomic DNA | OL147, OL148 | CYC1 terminator amplification |
| BRICK529 | *S. cerevisiae* genomic DNA | OL1747,<br>OL1748 | CYC1 terminator amplification |
| BRICK530 | *S. cerevisiae* genomic DNA | OL1749,<br>OL1750 | ADH1 terminator amplification |
| BRICK781 | pCfB8800 | OL2668,<br>OL2669 | yeGFP amplification |
| SYN92 |  |  | Synthetic DNA for CYC1 promoter,<br>sequence found in table 6. |

**Table 6.**
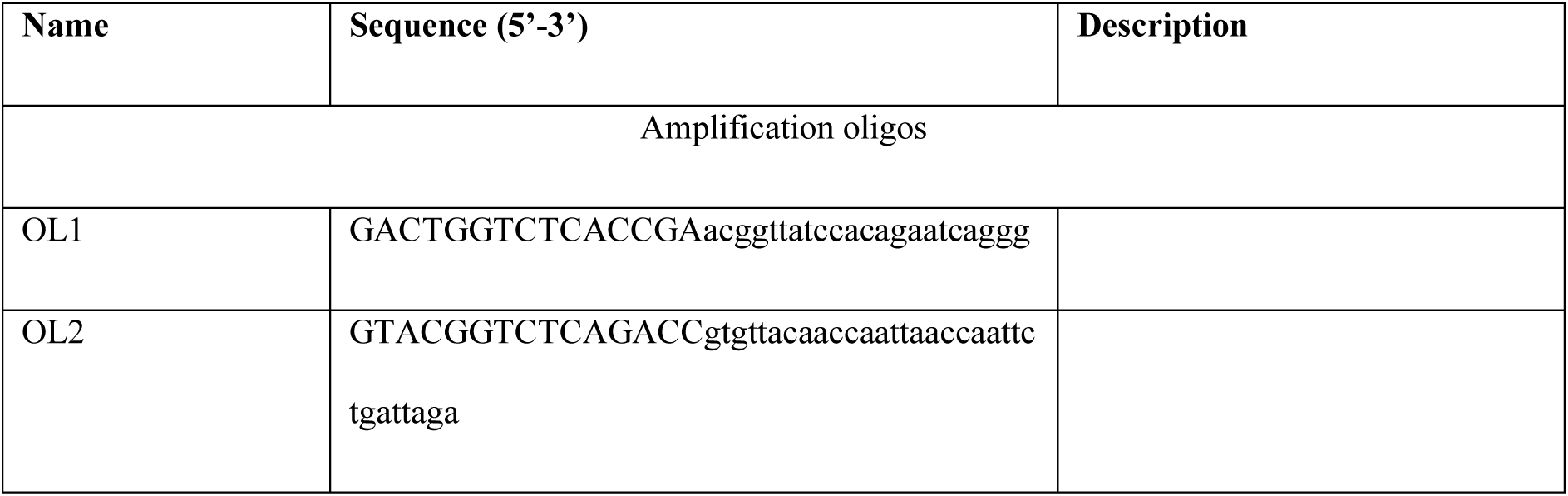
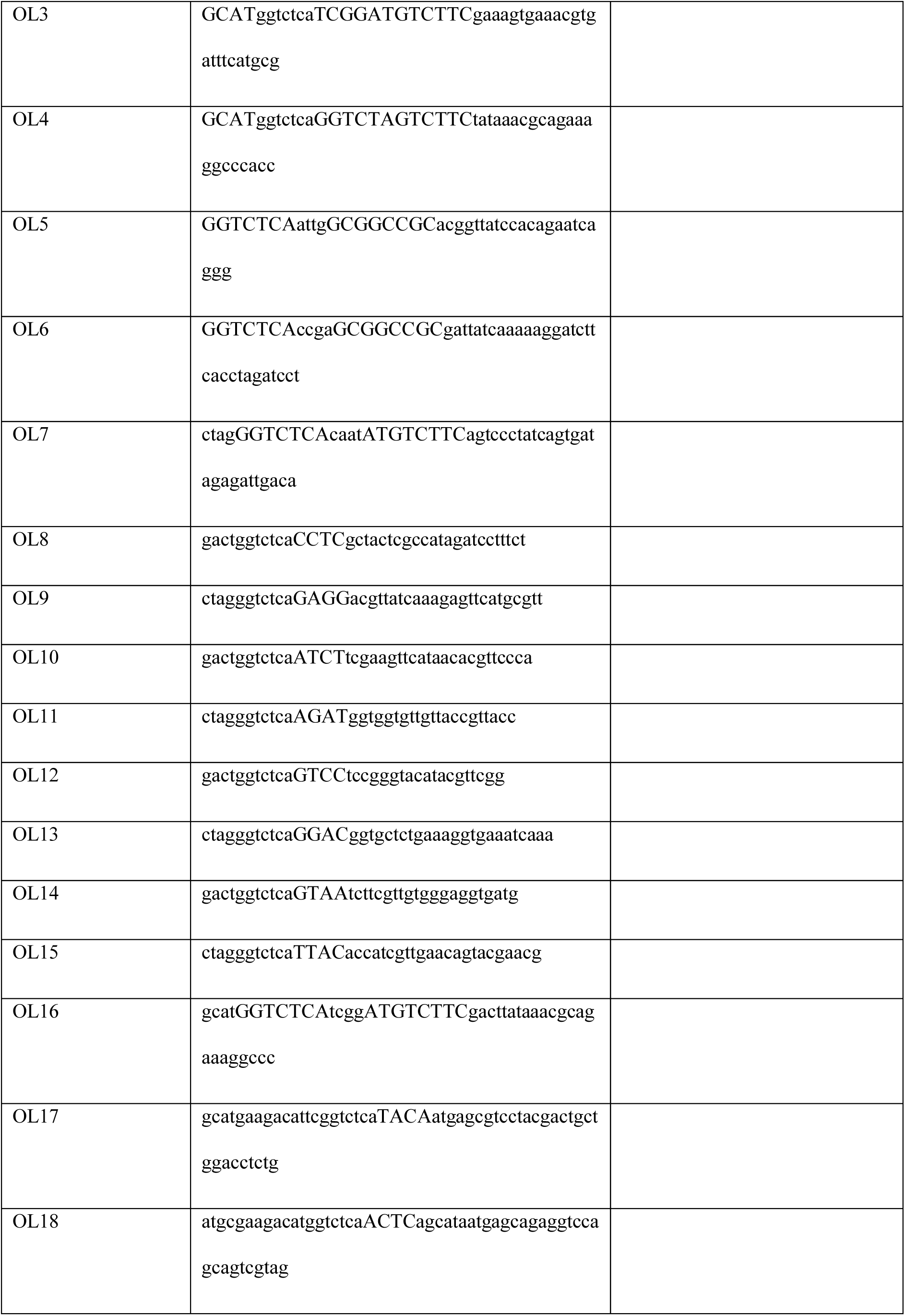
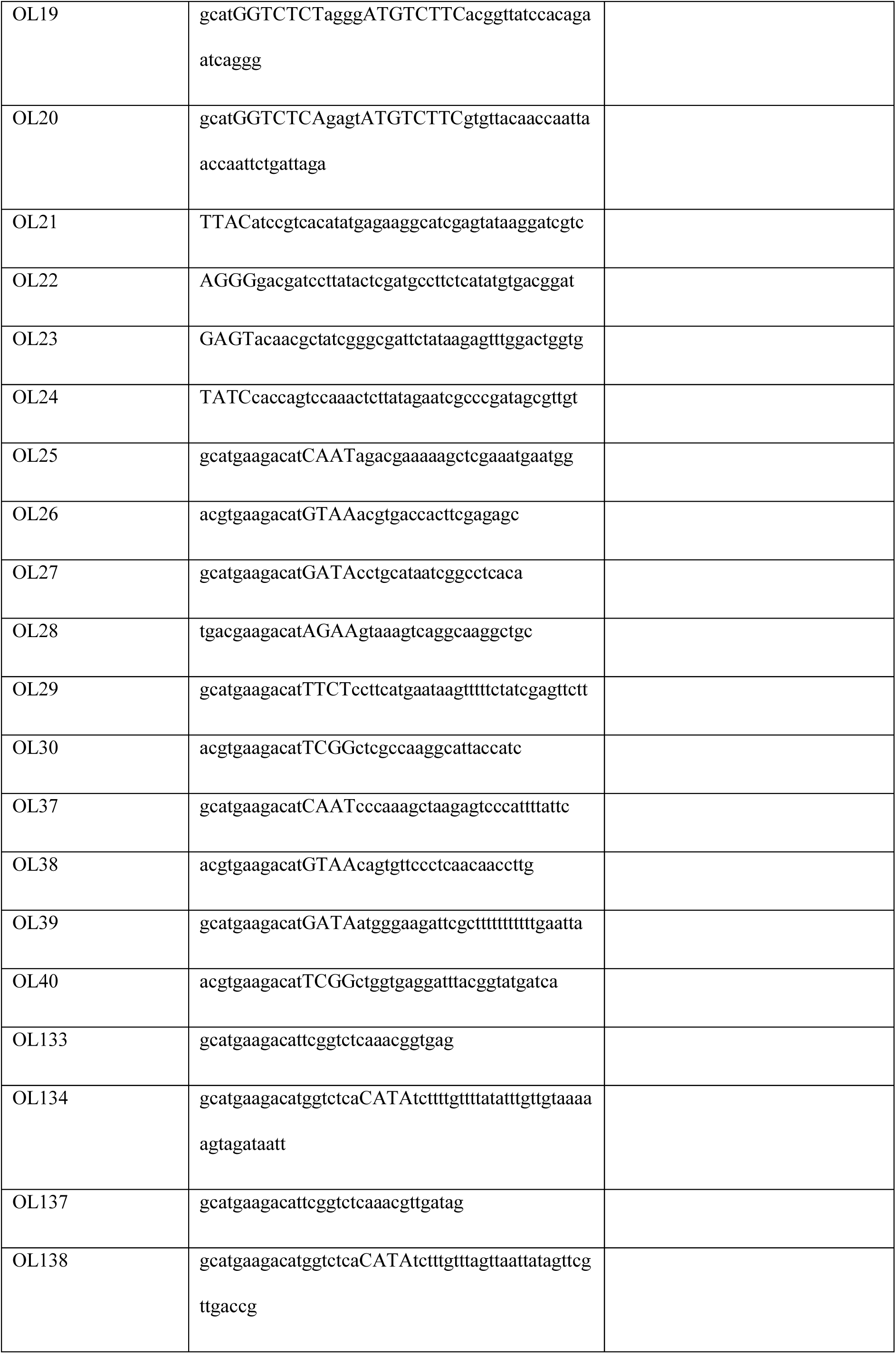
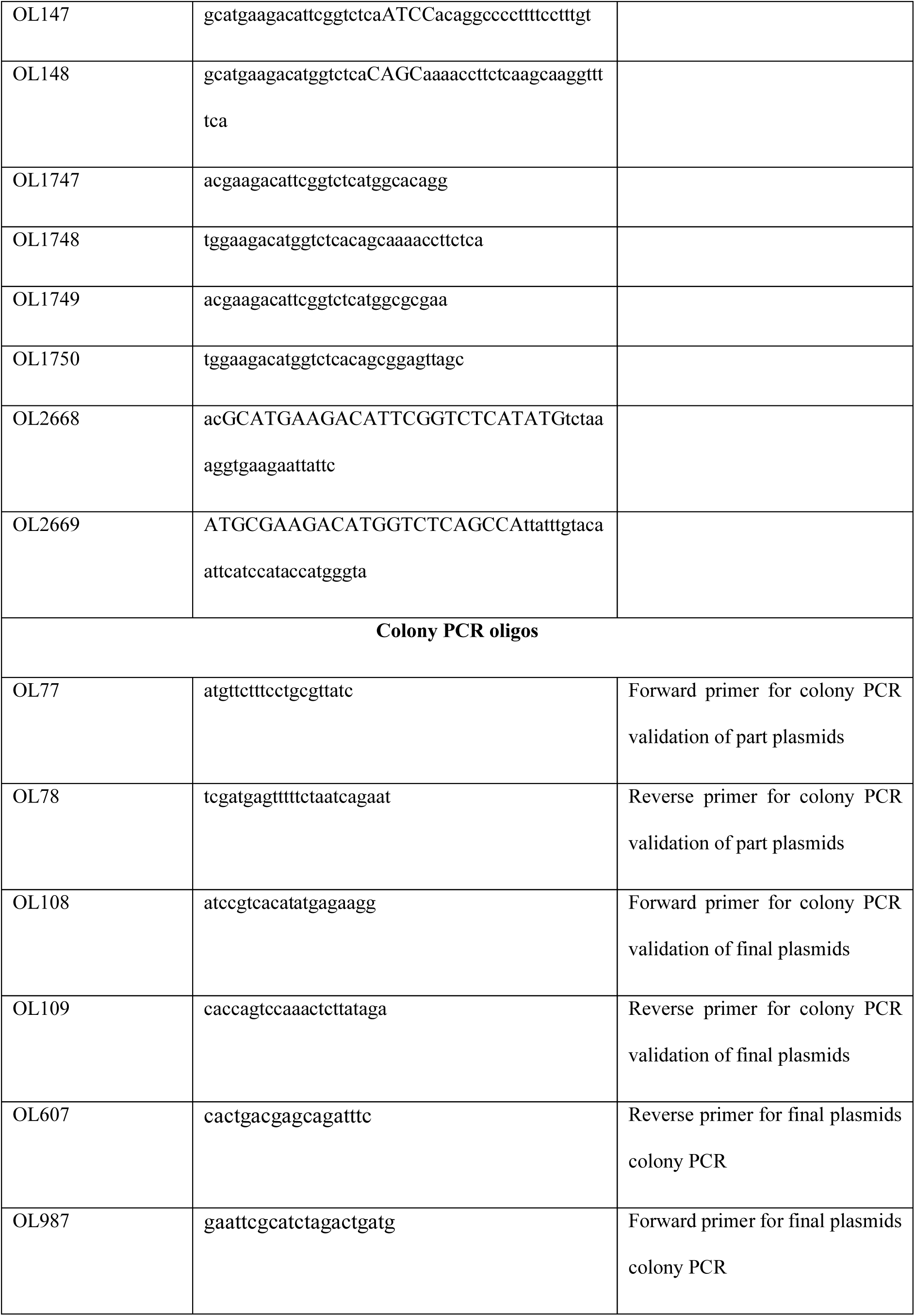
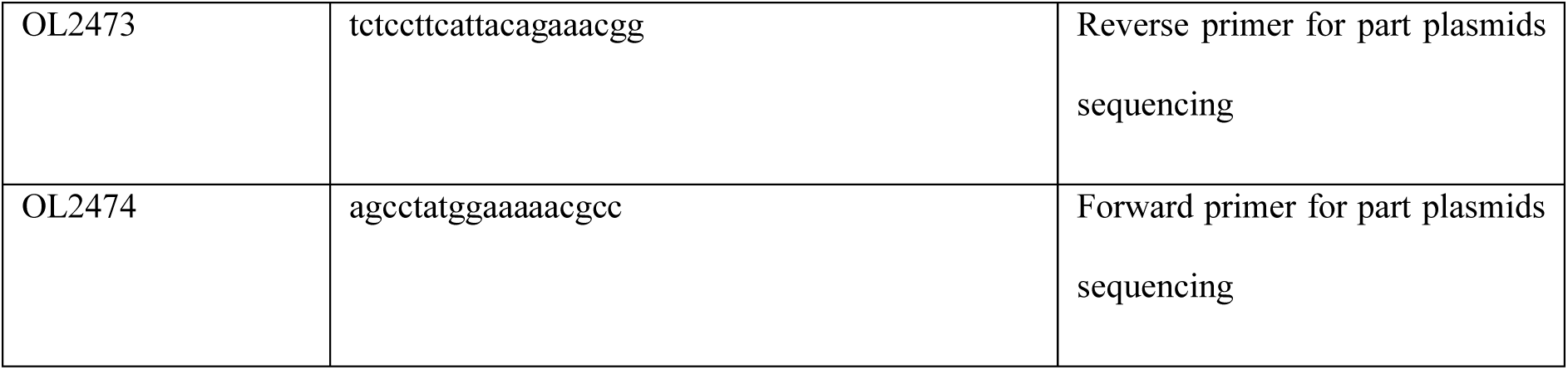
Oligos used in this study.

**Table 7.**
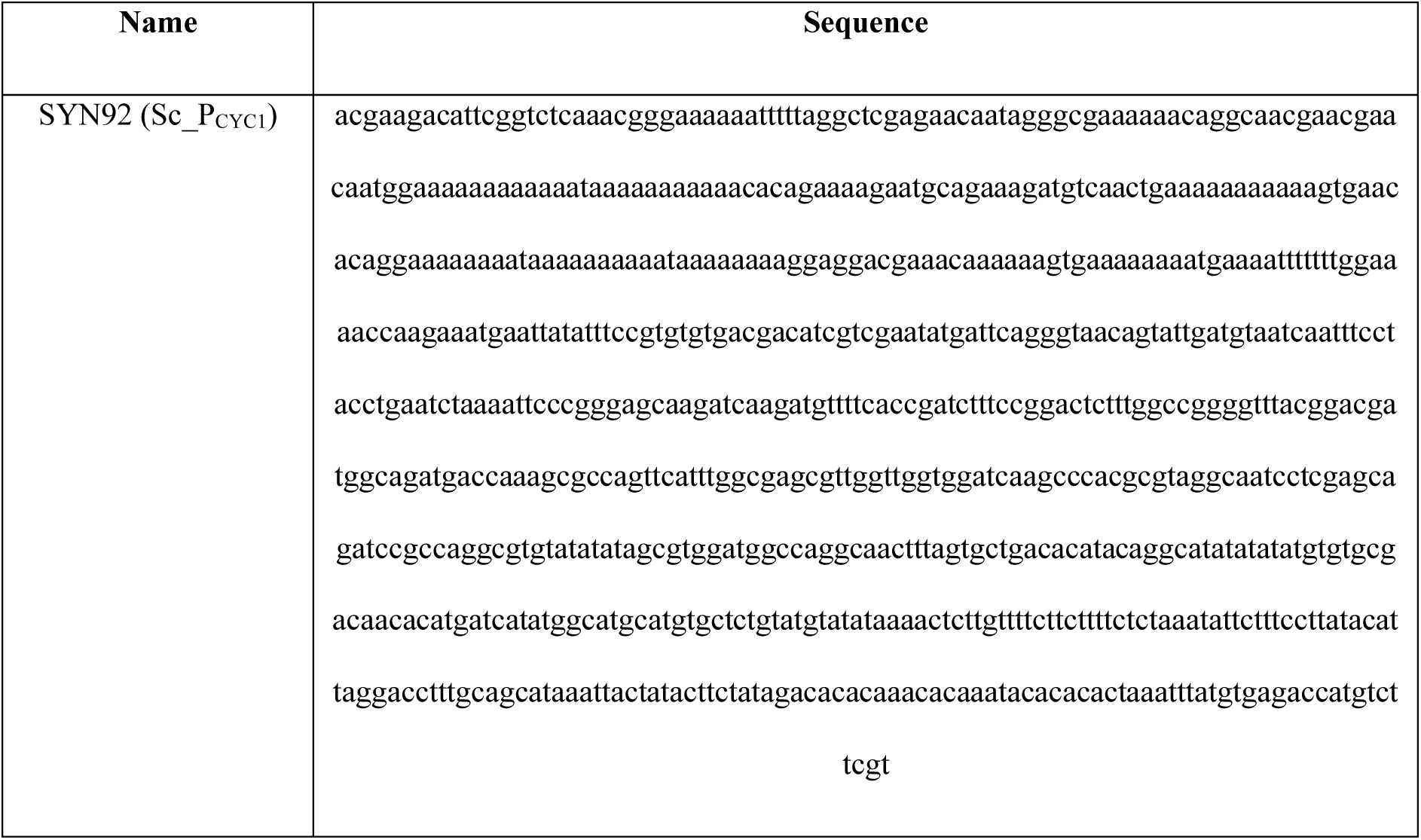
Synthetic DNA used in this study.

Assemblies were transformed into *E. coli* and selected on LB agar supplemented with 50 mg/L kanamycin (part plasmids) or 100 mg/L carbenicillin (final plasmids containing repair template). Colonies were screened via colony PCR (cPCR) using the FIREPol Master Mix (Solis BioDyne), and plasmids were isolated using FavorPrepTM Plasmid Extraction Mini Kit. All part plasmids were additionally validated through Sanger sequencing (University of Tartu – Core Facility of Genomics). To obtain repair dsDNA templates, final plasmids containing the GFP-expressing cassettes were linearized with FastDigest NotI (Thermo Scientific).

#### 2.2.3. Yeast transformation and strain confirmation

*S. cerevisiae* expressing SpCas9 was transformed with a linearized dsDNA repair template and a guide RNA expressing plasmid, following the lithium acetate/single-stranded carrier DNA/polyethylene glycol method [18]. Transformants were selected on YPD agar plates with 100 mg/L nourseothricin and evaluated for their GFP expression levels (Table 3). Genomic DNA was extracted using lithium acetate (LiOAc)/SDS [19] and correct integration was verified by cPCR (FIREPol Master Mix, Solis BioDyne).

#### 2.2.4. GFP expression analysis

Confirmed strains were grown overnight in CDM, diluted to a starting OD600 of 0.02 in fresh CDM media and transferred to microtiter plates. Growth (OD600) and GFP fluorescence (Ex 479 nm, Em 520 nm) were monitored every 10 minutes for 48 h at 30°C using a BioTek Synergy H1 microplate reader (Figure 4).

## 3. Results

### 3.1. Sensitivity evaluation using dye-filled droplets

To evaluate the performance of the sorters optical detection system, a FITC dilution series was used to characterize sensor response and estimate the effective limit of detection. Droplets containing defined FITC concentrations were measured at two digital-to-analogue converter (DAC) (similar to gain) settings, 0,65 V and 1,25 V (Figure 3).

**Figure 3.**
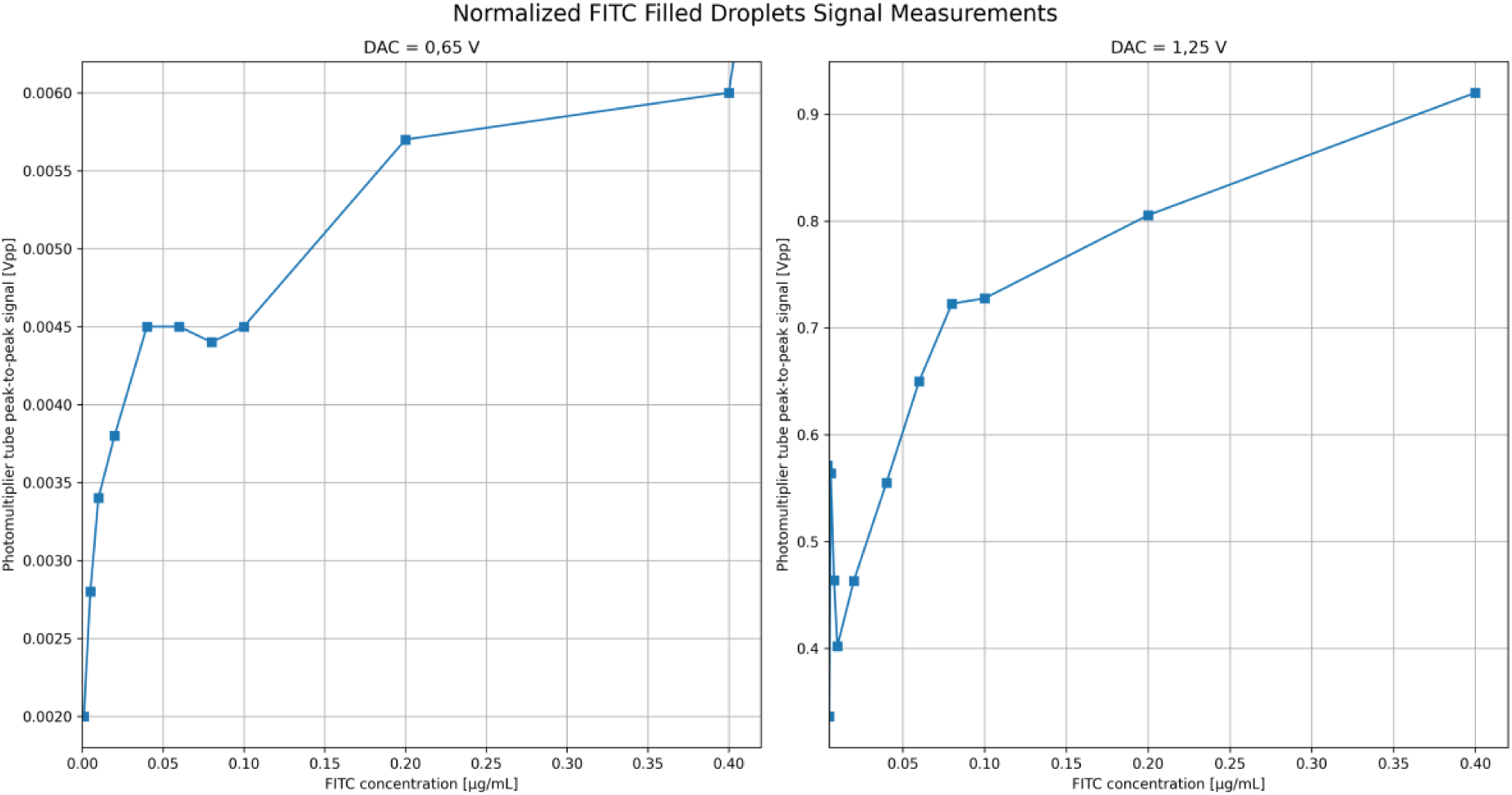
Normalized FITC-filled droplet signal versus FITC concentration at two DAC voltage settings (0,65 V and 1,25 V). The DAC (digital-to-analogue converter) voltage controls the analogue signal level in the detection system affecting output amplitude and signal-to-noise ratio, similar to gain in microscopy.

At very low concentrations (≤0,001 µg/mL), signal variability and deviations from linearity were observed. These effects likely arise due to preparation noise and droplet-to-droplet variation in the ∼120 pL droplets. At the lower DAC setting (0,65 V), the system was capable of detecting low fluorescence signals, although distinguishing signal from background noise was challenging. Increasing the DAC value reduced signal variability and improved signal separation across a broader concentration range. At 1,25 V, the measured signal showed a generally increasing trend with FITC concentration over a broader range, although minor fluctuations remained at very low signal levels.

Overall, the results confirm that the system can reliably detect fluorescence differences across a range of concentrations, however with a trade-off between sensitivity and dynamic range. Lower DAC values extend the measurable range but reduce signal-to-noise ratio, while higher DAC values improve signal differentiation but reduce the dynamic range due to the measurement ceiling of the detection electronics.

### 3.2. Droplet-based fluorescence analysis

Fluorescence measurements of droplets containing yeast cells revealed broad signal distributions arising from both biological and technical variability. Factors such as the number of cells per droplet, cellular growth state and intrinsic fluorescence contributed to multimodal intensity profiles (Figure 4).

**Figure 4.**
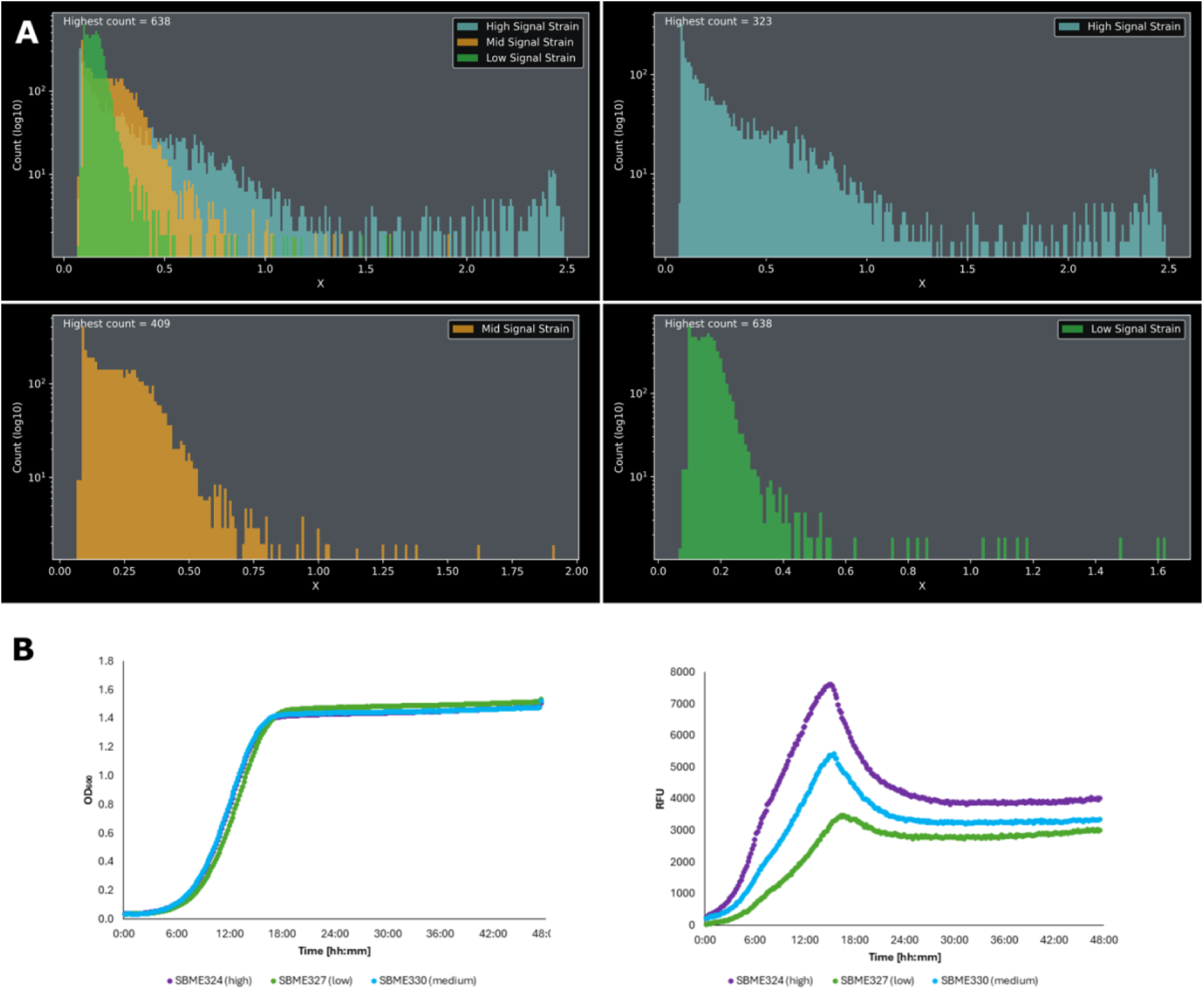
Signal strength distributions for high-, mid- and low-signal yeast strains (Table 3). (**A**) Fluorescence intensity histograms of the three strains together with an overlapped histogram. The x-axis represents the maximum voltage peak of individual droplets, while the y-axis shows the droplet count (number of droplets passing through the detector with a given signal), displayed on a logarithmic scale. The graph illustrates the distribution of fluorescence signals from droplets analysed by the detector. Droplets consist of liquid yeast medium containing suspended yeast cells with a theoretical fill rate of 66%. Empty droplets contain only yeast medium, which exhibits background fluorescence and produces a signal below 0,2. (**B**) Microtiter plate readings for the *S. cerevisiae* strains used in fluorescence-assisted droplet sorting. OD_600_ readings indicate biomass growth over time, while relative fluorescence unit (RFU) measurements show GFP fluorescence levels of the strains, confirming their different fluorescence intensities measured with a microtiter plate reader.

To better interpret these signal distributions, fluorescence intensity windows were analysed using high-speed image capture rather than active sorting. This enabled correlation between measured fluorescence signals and droplet contents. Analysis indicated that high intensity signals could result from droplets containing multiple cells, increased per cell fluorescence or a combination of both factors.

When analysing droplet-in-oil samples containing individual strains, the resulting signal distributions were sufficiently distinct to allow separation between empty droplets and droplets containing cells. However, when droplet-in-oil samples containing strains were combined, significant overlap between fluorescence distributions was observed. While extreme populations (for example empty droplets vs highly fluorescent droplets) remained distinguishable, separation between strains with similar fluorescence levels was less reliable.

These observations highlight a limitation of single parameter fluorescence-based selection and motivate careful threshold selection during sorting experiments.

### 3.3. Proof-of-concept sorting

To demonstrate system functionality, proof-of-concept sorting experiments were performed using droplets containing yeast cells (Figure 5). Sorting was triggered based on fluorescence thresholds derived from the prior characterization.

**Figure 5.**
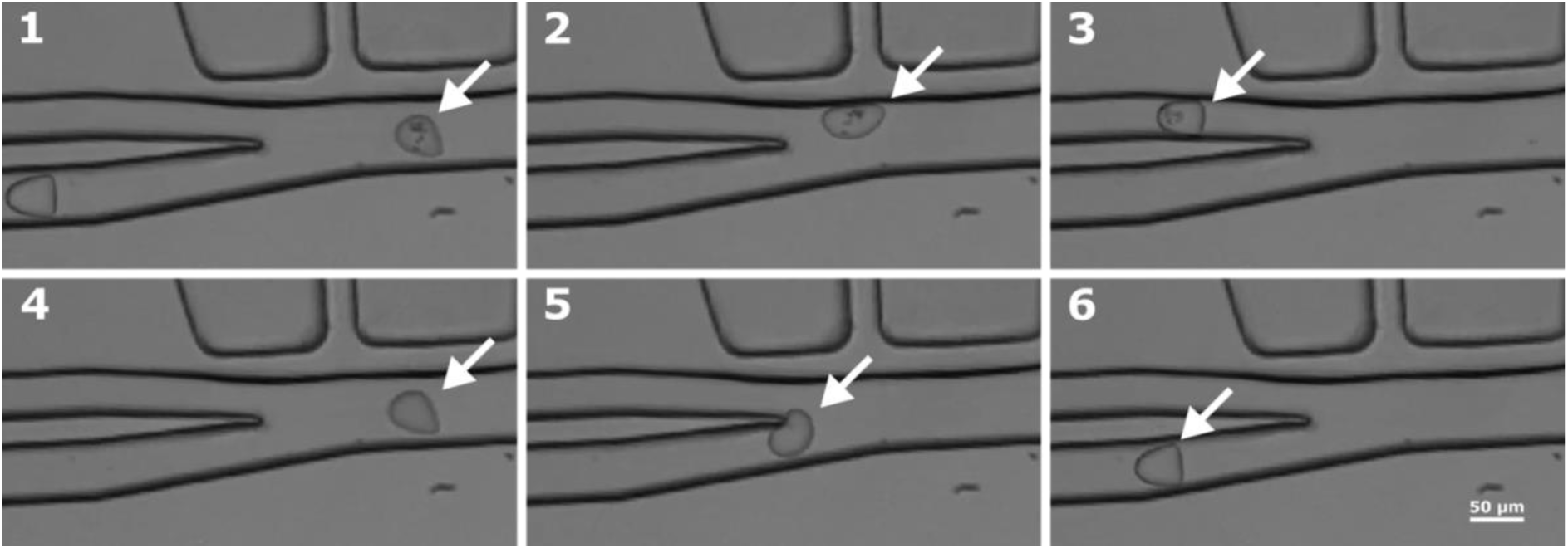
Snapshots of droplets during sorting when the electrode is triggered after the fluorescence signal exceeds the predefined detection threshold, as measured by optical fibres. Panels (1-3) show the sorting process for droplets that exceed the threshold. Once detected by the optical fibres, droplets are initially directed toward the waste (lower) channel by default. After a predefined delay, a current is applied to the sorting electrodes, generating a dielectrophoretic force within the droplet. This force pulls the droplet toward the electrodes, causing it to divert and exit through the sorted (upper) channel. Panels (5-8) show droplets that do not exceed the threshold. In this case, the sorting electrodes are not activated, no dielectrophoretic force is generated, and the droplets continue along their default path, exiting through the waste (lower) channel.

The system successfully redirected target droplets into the collection outlet when the timing between fluorescence detection and redirection was correctly adjusted and sufficient dielectrophoretic force was applied. Because sorting parameters such as pulse timing, duration and voltage are tuneable, operating conditions could be achieved in which the sorted fraction showed enrichment of fluorescent droplets compared with both the waste fraction and the original input sample.

Although sorting accuracy was limited by overlapping signal distributions in mixed strain samples, these experiments demonstrate successful end-to-end operation of the platform, from optical detection to droplet redirection.

## 4. Conclusion

This work presents an early technical preview of the DropletFactory CORE FADS platform. Sensitivity measurements using FITC containing droplets demonstrated reliable fluorescence detection and highlighted the relationship between detector gain, sensitivity and dynamic range.

Analysis of GFP expressing yeast droplets revealed expected signal variability due to biological and technical factors. Proof-of-concept experiments further confirmed successful droplet sorting based on fluorescence thresholds, demonstrating end-toe-end functionality of the system

## 5. Acknowledgements

We thank Dr. Tomasz Kaminski (University of Warsaw) for continuous awesome advice and providing the microfluidic chips. This project has received funding from the Pathfinder Open under the European Union’s Horizon EU research and innovation programme, project 3D-BRICKS (Grant agreement No. 101099125). This work was (partially) supported by the ASTRA+ programme “Increasing the impact of research and strengthening the knowledge transfer capacity of research organisations and higher education institutions”, co-funded by the European Union (project No. 2021-2027.1.01.25-1163 and 2021-2027.1.01.25-1019).

## 7. Appendix

## Notes

### Competing Interest Statement

The authors have declared no competing interest.

